# The Work Performed By Biological Motors

**DOI:** 10.1101/580241

**Authors:** Josh E. Baker

## Abstract

Molecular motors are enzymes that perform work (*F* · *x*) when they move along a track a distance *x* against a constant force *F*. This work is performed through intermediate chemical steps in a motor’s ATPase reaction cycle, each step having a free energy change associated with it that is a sum of chemical, Δµ_chem_, and mechanical, Δµ_ext_, potentials. Defining Δµ_ext_ is fundamental to our understanding of how molecular motors work, yet after decades of study the definition of Δµ_ext_ remains disputed. Some postulate that Δµ_ext_ is a function of both *F* and *x*, while others assume that Δµ_ext_ is a function of neither *F* nor *x*, and still others argue that Δµ_ext_ is a function of *F* but not *x*. Here we evaluate these models and conclude that only the latter – a mechanochemical model proposed by A.V. Hill in the 1930’s – describes molecular motor mechanochemistry.

## Main Text

Intermediate chemical steps in a motor-catalyzed ATPase reaction can generate force and movement along tracks (*1*–*5*). Figure 1 illustrates three unique molecular mechanisms (top) having different sequential relationships between motor chemistry, movement, and force generation (middle) with corresponding free energy landscapes (bottom). In 1957, A.F. Huxley proposed a model for molecular motors (Fig. 1, Captured Chemical Strain), which was subsequently formalized by T.L. Hill (*6*, *7*) in which a motor generates force (Strain) when thermal fluctuations in the displacement, *d*, of a molecular spring are captured in a strained state upon motor-track binding (Chemistry), and the motor moves a distance, *x*, (Movement) only upon relaxation of the molecular spring in what is referred to as a powerstroke. In 1974, Huxley and Simmons presented a modification of this model (Fig. 1, Chemical Strain) in which a motor, when bound to a track, undergoes a chemical working step (a motor rotation) that stretches a molecular spring a distance *d* (Chemistry-Strain), generating mechanical strain and force (*7*, *8*). Consistent with the Hill formalism, in this model a motor moves a distance, *x*, only upon relaxation of the molecular spring (Strain-Movement). In the 1990’s, direct measurements of the distance, *x*, a motor moves against an apparent constant force, *F*, (*9*, *10*) inspired the development models (Fig. 1, Chemical Fx) whereby movement and thus Fx-work are directly performed with a motor’s working chemical step (Chemistry-Movement).

**Figure 1.**
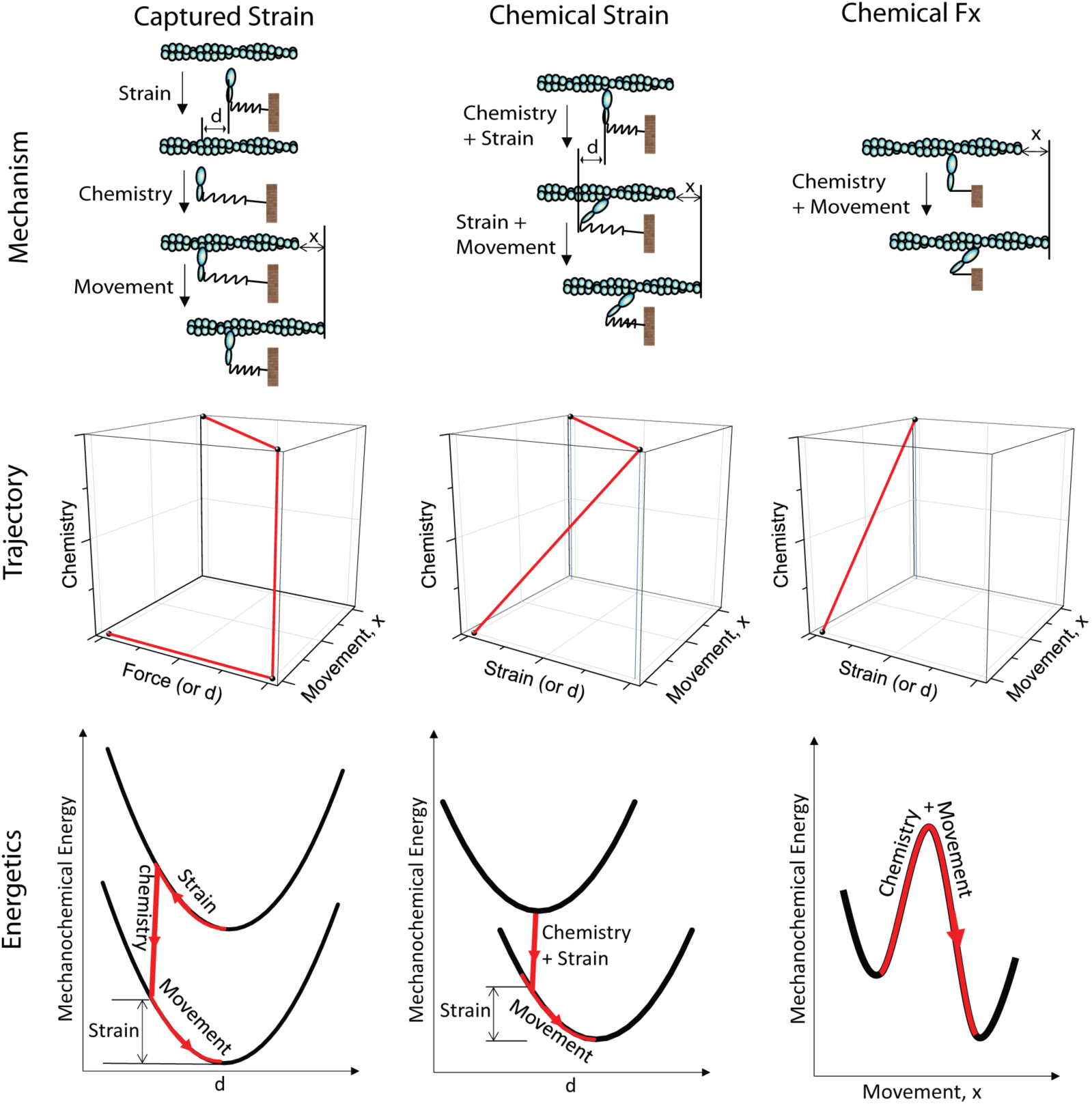
Three proposed models describing different sequential relationships between motor force generation (Strain), motor movement, and motor chemistry. Mechanisms depict motors as ovals, tracks as a string of spheres, motor strain/force generation as a molecular spring stretched a distance, *d*, and movement as track movement through a distance, *x*. Trajectories are a graphical representation of the sequence of force generation, movement and chemistry proposed for each model. Energetics show the paths through the energy landscapes that proposed for each model.

To determine which of these models is correct we focus on their contrasting definitions of mechanical potentials, Δµ_ext_. In general, for biological motors, Δµ_ext_ is the reversible mechanical work performed with a motor’s working chemical step. It is reversible because it is energy that is part of (contained in, available for use by, and measurable within) the motor system. It is important to distinguish between this reversible work and the irreversible, or dissipative, work performed by molecular motors. The chemical reactions that drive motor mechanics are exothermic, which means that motor catalysts lose heat irreversibly from the motor system to their surroundings. For example, the work performed by myosin motors in our legs when we climb a hill lifting our bodies against gravitational forces is irreversible work because it cannot be recovered by the myosin motors in our legs on the way back down. This is because the net heat lost from our bodies with the exothermic chemical reaction that fueled our uphill work cannot be macroscopically reabsorbed by our bodies to reverse motor working steps on the way down. In contrast, a reversible mechanical potential, Δµ_ext_, can be used to reverse motor chemistry.

The above models differ fundamentally in their descriptions of Δµ_ext_ (Fig. 2) In chemical Fx models Δµ_ext_ equals *F·x* (Fig. 2A). In contrast, in chemical strain models Δµ_ext_ is not a function of *x* but – depending on specific assumptions described below – may or may not be a function of *F* (Figs. 2B and 2C). This lack of clarity on motor fundamentals must be resolved before we can ever claim to understand how molecular motors work.

**Fig. 2.**
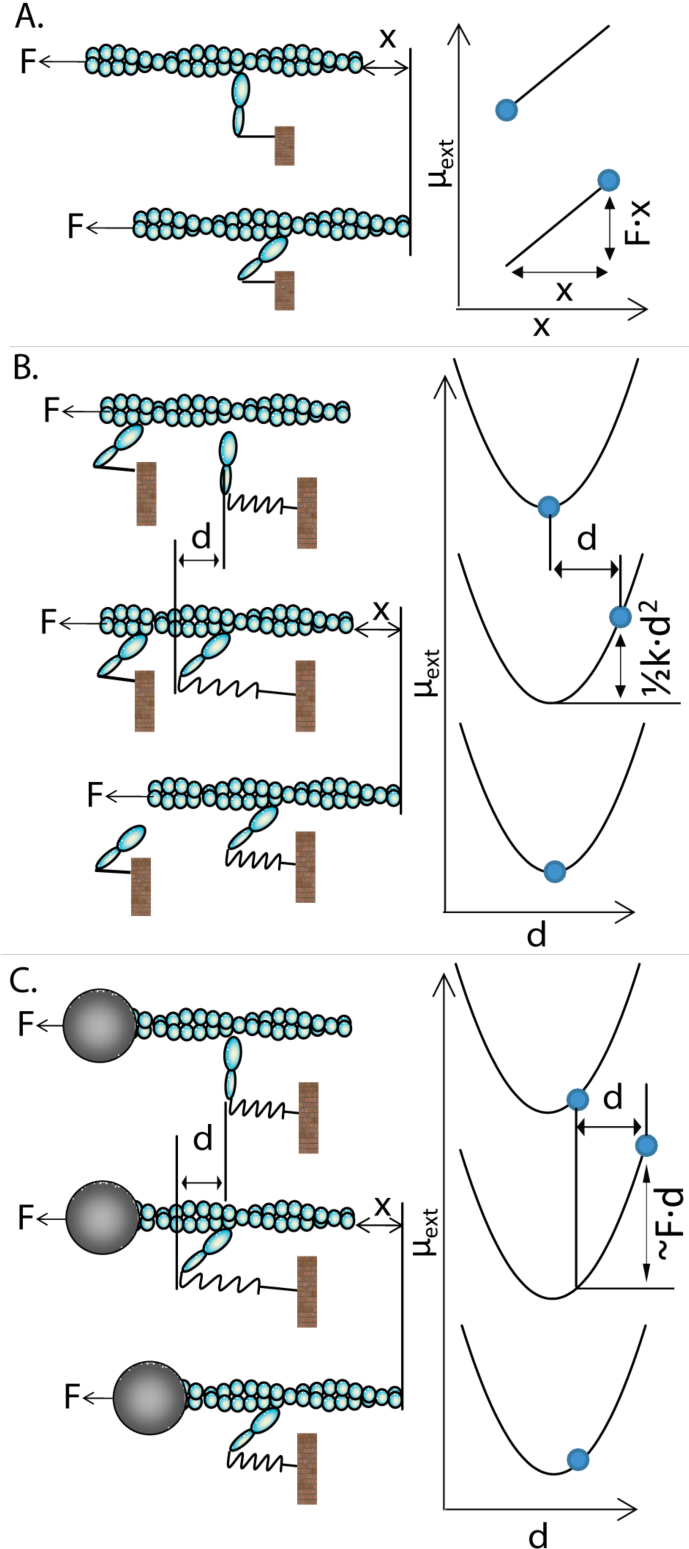
Three disparate models describing mechanochemical coupling in motors that move along a track a distance, *x*, against a constant force, *F*. Left panels show structural mechanisms. Right panels show corresponding mechanical potentials. (A) According to chemical Fx models, a motor working step (the rotation of a motor [double ovals] bound to a track [string of spheres]) moves a track a distance, *x*, against a constant force, *F*, performing reversible work, Δµ_ext_ *= F·x*. (B) According to Huxley-Hill chemical strain models, a mechanical barrier (motor on left) prevents the working step from moving the track against a constant force, *F*. Instead, the working step stretches local spring-like elements (spring), generating force and performing work, Δµ_ext_ = ½*k*_*uni*_·*d*^2^ (top to middle), which is a function of neither *F* nor *x*. With the release of the mechanical barrier (middle to bottom), movement, *x*, occurs with spring relaxation and Δµ_ext_ is irreversibly lost as Fx work. (C) Equilibrated chemical strain models resemble Huxley-Hill except the barrier is replaced with a damping element (large sphere), resulting in a mechanical potential, Δµ_ext_ ≈ *F·d*, that is a function of *F* but not *x*.

In chemical Fx models, *F* is assumed to be a constant “barometric” force, and *x* a reaction coordinate (*9*) (Fig. 1, Chemical Fx). According to this model, when a motor moves along its track a distance, *x*, against a constant force, *F*, the Fx-work performed is the mechanical potential, Δµ_ext_ = *F·x* (Fig. 2A, top to bottom), which is assumed to be reversible (*9*, *11*). However, it is easy to show that Fx-work is not a reversible mechanical potential but rather irreversible work that is lost from the motor system. Figure 2 illustrates a processive motor such as kinesin or myosin V (*12*–*14*) that performs Fx-work when it walks around a circle against a force, *F*, exerted parallel to the track. Held at a constant *F*, the motor begins and ends in the exact same mechanical state, which means that even though net Fx work was performed, that work is not physically contained, available for use, or measurable within the motor system. Instead, Fx work is lost from the motor system because it is performed with a macroscopically irreversible exothermic chemical reaction. It has been argued that chemical Fx-work models are consistent with observed motor stepping in single molecule studies (*15*–*17*); however a careful analysis of these experiments (Fig. 3) shows that what has been assumed to be reversible Fx-work in these experiments is in fact irreversible work. Consistent with Huxley-Hill, this analysis shows that the basic mechanism by which Fx-work is lost from a motor system is the motor powerstroke.

**Figure 3.**
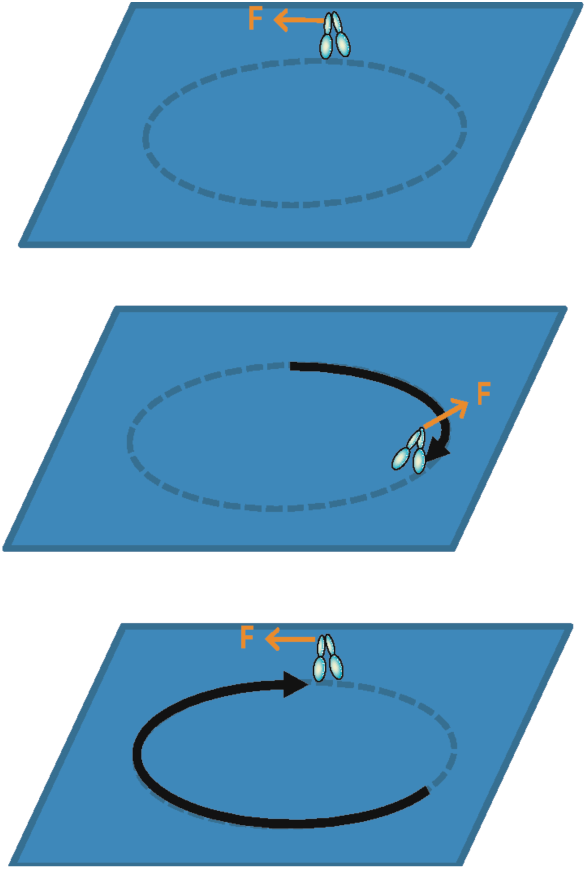
A motor (illustrated as a two-headed processive motor) moves a distance *X* around a circular track against a constant force, *F*, applied tangent to the track. Each dash on the track represents a single motor step, *x*, associated with a single motor reaction cycle. According to Fx work models, the mechanical potential of the motor system increases by *Fx* around each chemical cycle and by *FX* each time the motor circles the track, when in fact the state of the motor system is the same at the beginning (top) as it is at the end (bottom).

Figure 3A illustrates a typical single motor mechanics experiment in which a motor displaces a bead held in an optical trap. The trapped bead behaves like a linear spring with stiffness *κ*_*trap*_, such that when the bead is displaced a distance *d* from the trap center (Fig. 3A, top to bottom) strain energy, ½*κ*_*trap*_·*d*^2^, and force, κ_trap_·d, are generated. Here, for simplicity, we assume that *κ*_*trap*_ is much less than the stiffness of the motor. In this experiment, consistent with chemical strain models (Figs. 1C), Δµ_ext_ (= ½*κ*_*trap*_·*d*^2^) is the strain generated with the chemical motor’s working step.

Figure 3B illustrates a similar experiment, only with the optical trap held at a “constant force”. Here, the trap spring is initially held at a constant force, *F*, by positioning the optical trap at a computer-controlled displacement of *F*/*κ*_*trap*_. When a motor working step displaces the trap spring an additional distance, *d*, (Fig. 3B, top to middle) the computer-controlled feedback system responds to undo the force generated with this displacement by moving the trap center a distance *x* (= *d*) (Fig. 3B, middle to bottom).

In this experiment, the mechanical work, Δµ_ext_, performed by the motor working step is chemical-strain work (Fig. 3B, top to middle, Fig. 2A) not chemical-Fx work (Fig. 2B). Specifically, the strain generated in stretching the trap spring a distance *d* from an initial position *F*/*κ*_*trap*_ is the difference between the final, ½*κ*_*trap*_(*F*/*κ*_*trap*_ + *d*)^2^, and initial, ½*κ*_*trap*_(*F*/*κ*_*trap*_)^2^, strain, or

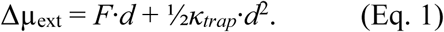

Movement, x, (i.e., Fx work) occurs with the subsequent feedback response (Fig. 4B, middle to bottom), but this movement is not reversible; it is dissipative. When the trap moves a distance, x, the work done with the working step in stretching the trap, Δµ_ext_, is lost from the motor system (the trap spring) to the computer-controlled stepper motor (SM) that moved the trap, resulting in no net change in the mechanical state (Δµ_ext_) of the motor system for the overall two-step transition (Fig. 4B, top to bottom). In other words, similar to Fig. 2, the mechanical state of the motor system is the same after the step at is was before. This mechanochemistry is consistent with chemical-strain (Fig. 2A) not chemical-Fx models (Fig. 2B), with a working step generating strain energy and the irreversible relaxation of this strain through what is essentially a powerstroke mechanism performing Fx-work. In these experiments, a motor’s working step need not satisfy detailed balance, since the computer algorithm that controls the SM is not programmed to conserve energy. In fact, it can be shown that in these experiments when a forward step is followed by a reverse step, a molecular motor dissipates net mechanical energy, *κ*_*trap*_·*d*^2^, to the SM.

**Fig. 4.**
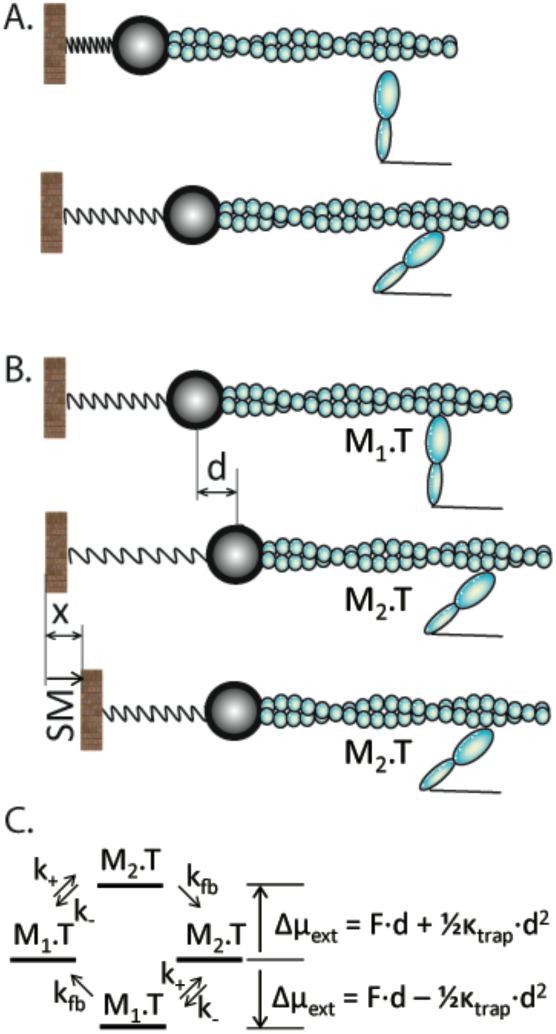
Mechanochemical coupling in single motor optical trap experiments. Displacement of a bead (large circle) from the optical trap center (rectangle) has a force response similar to that of a linear spring. (A) Upon track binding, a motor’s chemical working step displaces the track and associated bead a distance, *d*, generating a mechanical strain, Δµ_ext_ = ½*k*_*trap*_·*d*^2^, in a trap spring of stiffness *k*_*trap*_. (B) In a force-feedback optical trap, the trap is held at a pseudo-constant force via a feedback system. When a motor’s mechanochemical step (from M_1_.T to M_2_.T) stretches the trap spring a distance *d*, a stepper motor (SM) responds by moving the trap a distance *x* (= *d*), maintaining a constant force. (C) A kinetic scheme for the reversible two-step processes illustrated in (B). A motor’s mechanochemical step (M_1_.T to M_2_.T) is reversible and has forward and reverse rate constants of *k*_*+*_ and *k*_-_, respectively. The feedback response occurs on a time scale of 1/*k*_*fb*_. The change in mechanical strain, Δµ_ext_, with an M_1_.T to M_2_.T transition is greater than the mechanical strain dissipated with an M_2_ to M_1_ transition.

The kinetics and energetics for the working step in this “constant force” assay (Fig. 4B) are illustrated in Fig. 3C. The apparent rates for the forward and reverse steps are *k*_*app+*_ = *k*_*+*_·*k*_*fb*_/(*k*_-_ + *k*_*fb*_) and *k*_*app-*_ = *k*_-_·*k*_*fb*_/(*k*_*+*_ + *k*_*fb*_), respectively, where *k*_*+*_ and *k*_-_ are the forward and reverse rates for the motor chemical transition and k_fb_ is the inverse dead time for the feedback response. Because *k*_*fb*_ is typically much faster than either *k*_-_ or *k*_*+*_, it is often assumed that *k*_*app+*_ = *k*_*+*_ and *k*_*app-*_ = *k*_-_. According to Fig. 4B, the strain-dependences of these rates are

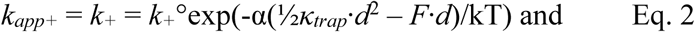

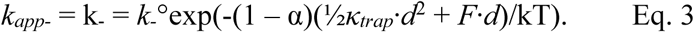

where *k*_*+*_° and *k*_-_° are rate constants at zero strain, and α is the fraction of Δµ_ext_ generated prior to the activation energy barrier for the forward chemical step. Thus even though no net mechanical energy is generated with the overall working step, *k*_*app+*_ and *k*_*app-*_ are strain-dependent because strain is transiently generated with the rate limiting steps for these transitions. The reason that Fx-dependent kinetics (Fig. 2B) accurately describes the kinetics of motor stepping in these single molecule studies is that when *d* is small relative to *F*/*κ*_*trap*_ (which is typically the case), Eq. 1 becomes

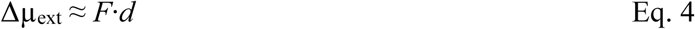

which in this particular example (but not always) equals *F*·*x* because the distance the SM moves the trap, *x*, is equal to the displacement generated by the motor working step.

Even if we could engineer an infinitely fast detection and force feedback system, Fx-work would still be dissipated in these experiments since the thermal fluctuations of a motor that activate a working step and generate thermal forces in the direction of movement would be dissipated immediately by the rapid force feedback that maintains a constant force. In effect a motor’s working step would be thermally cooled by the rapid force feedback and thus would never occur.

To allow thermally-activated motors to move along *x*, the chemical forces of molecular motors must either be uncoupled from *F* or be coupled to a dynamic external force, *F(t*), that becomes part of the motor’s energy landscape and chemistry. The former is essentially the “independent force generation” by motors assumed in Huxley-Hill chemical-strain models. The latter is the collective force generator model we proposed in 1999, which is consistent with A.V. Hill’s macroscopic model of muscle energetics.

In Huxley-Hill models, chemical forces of individual motors are allowed to fluctuate because they are assumed to be mechanically isolated (uncoupled) from a constant external *F* (Fig. 2B). This is consistent with an Eyring-like model where mechanical strain is first generated locally and then subsequently used to generate global movement, *x*. The effective mechanical barrier (Fig. 2B) that isolates motor mechanics (molecular springs) from global mechanics, *F*, is assumed to result from over-damping of movement, *x*. Here a motor’s working step does not directly generate movement, *x*, and thus *x* does not contribute to Δµ_ext_. Similarly, because the motor working step is isolated from the constant external force, *F, F* does not contribute to Δµ_ext_ (*18*, *19*). This is in sharp contrast to chemical Fx models where both *F* and *x* contribute to Δµ_ext_.

According to Huxley-Hill chemical-strain models, a motor’s chemical working step is only “locally equilibrated” (*19*), stretching molecular compliant elements (springs) to generate local strain, Δµ_ext_ = ½*κ*_*uni*_·*d*^2^ (Fig. 2B, top to middle), where *κ*_*uni*_ is the motor stiffness and *d* is the motor step size. Motor movement, *x*, occurs subsequently through a relaxation of the motor spring, which is a dissipative transition that performs Fx work against *F* (the powerstroke). Claims that these models violate detailed balance (*11*) are based on the assumption that Fx work is conserved when in fact the Fx work performed with a powerstroke is dissipative (*20*).

In 1935, based on energetic measurements of contracting muscle, A.V. Hill proposed a model in which the conserved mechanical potential, Δµ_ext_, in muscle is a function of muscle force, *F*, but not movement *x*. In 1999, we made the first direct measurements of mechanochemical coupling in isometric muscle, similarly showing that Δµ_ext_ = *F*·*d* is a function of muscle force, *F*, and the motor step size, *d*, but not movement, *x*. These studies are at odds with chemical strain models that assume a motor’s working step is mechanically isolated from an external force, *F*; instead, they suggest that motors thermally equilibrate with *F*.

Following these studies, investigators have proposed muscle models that retain the localized molecular mechanical potential (i.e., independent force generation) of the Huxley-Hill chemical strain model while assuming that motors are thermally equilibrated with an external force (*21*–*23*). Figure 2C illustrates the general approach. Here a motor bound to a track equilibrates with an external force, displacing the molecular spring of a motor in its pre-working step state. A damping element (illustrated with a large bead) prevents the subsequent motor working step from moving against the external force, *F*, and localizes the work, Δµ_ext_, to molecular strain generated by that motor (independent force generation). Here because the motor in both the pre- and post-working step states equilibrates with the external force, the mechanical potential, Δµ_ext_, is a function of *F* (Fig. 2C, right). Specifically, at high *F*, Δµ_ext_ is approximately *F·d* (Eq. 4) consistent with our observations of mechanochemical coupling in muscle.

The problem with these mechanically equilibrated independent force models is that the lifetime of most track-bound pre-working step states is relatively short (sub millisecond) (*24*, *25*), and it is unlikely that a motor equilibrates with an external force on this time scale and not on the time scale of the working step (also sub milliseconds) (*24*, *25*). In 2000, we proposed a model in which rather than functioning as independent force generators, motors function as collective force generators, directly generating movement, x, against a dynamic external load F(t), coupling the external load to the chemistry and kinetic of the motor working step itself (*18*). This model is supported by subsequent studies showing that the working steps of myosin motors directly generate movement, *x* (*20*, *24*, *26*, *27*), and most recently we have shown that a frictional load (Fig. 4C with F = 0) dramatically inhibits the kinetics of the working step of myosin motors. This observation is inconsistent with the model in Fig. 4C, which predicts that the chemistry of the working step (Δµ_ext_) does not depend on the extent to which movement, *x*, is damped.

Consistent with A.V. Hill’s model of muscle, our model assumes a common, macroscopic compliant element that multiple motors collectively stretch (Fig. 5). According to this model, motors sequentially displace a single system compliant element having an effective stiffness, *κ*_*sys*_. With a working step one motor displaces the system spring a distance *d* performing work, Δµ_ext_ = ½*κ*_*sys*_·*d*^2^. A second motor then displaces the system spring an additional distance *d* performing work, Δµ_ext_ = ½*κ*_*sys*_·(2*d*)^2^ - ½*κ*_*sys*_·*d*^2^. This collective stepping continues until the work, Δµ_ext_, performed by a motor – which at high spring forces, *F*, is approximately Δµ_ext_ ≈ *F·d* (Eq. 4) – equals the free energy available for work with the working step, –ΔG, at which point force generation stalls at a force, *F*_*o*_ = –ΔG/*d*.

**Fig. 5.**
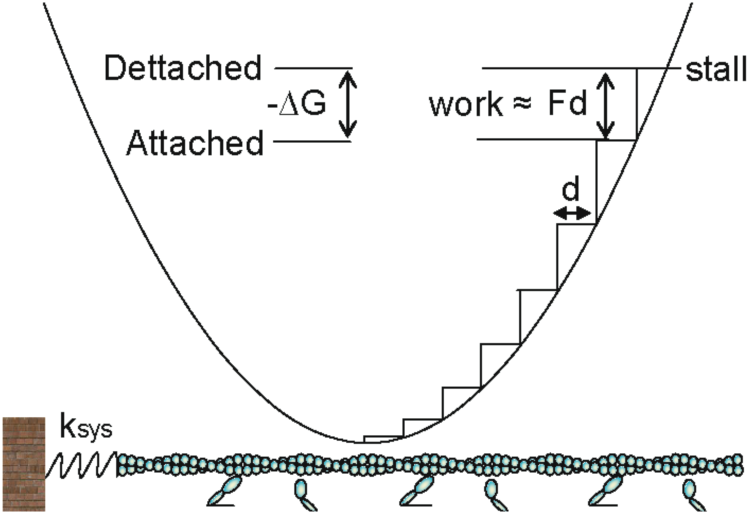
Collective force generation model. Motor working steps collectively stretch a single compliant element with stiffness *k*_*sys*_ (bottom) performing work (top) on the spring. With each sequential displacement, d, of the system spring, the energy and force, F, of the spring increases (top). At high F, a motor working step performs work, Δµ_ext_ ≈ *F·d*. With each subsequent step the spring force, *F*, and Δµ_ext_ = *F·d* increases until Δµ_ext_ = *F·d* equals the chemical free energy for the working step, -ΔG, at which point force generation stalls.

At forces, *F*, less than the stall force, *F*_*o*_ (= –ΔG/*d*), the total chemical free energy available for *Fx*-work is

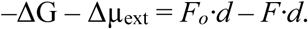

Conservation of energy mandates that the total chemical free energy equals the energy output by motors as work, *F·x*, and heat, *q*, or

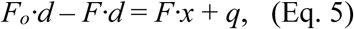

Again, we emphasize that the Fx-work performed by motors (*F*·*x* on the right side of Eq. 5) and the motor mechanical potential (Δµ_ext_ ≈ *F·d* on the left side of Eq. 5) are fundamentally different forms of work. The latter is conserved energy that is stored in the motor system as mechanical strain; the former is dissipated from the motor system with a motor powerstroke. To illustrate the difference, according to Eq. 5, as force, *F*, is collectively generated by motors, Δµ_ext_ ≈ *F·d* increases and *F*·*x* decreases until at stall force, Δµ_ext_ ≈ *F*_*o*_*·d* is at its maximum and *F*·*x* = 0.

Equation 4 is the energetic basis for the muscle F-V relationship that A.V. Hill proposed in 1938 (*18*, *28*). Indeed one could argue that, although they knew nothing of molecular motors, early muscle physiologists like Hill and Fenn understood the main conclusions of this paper. Hill understood that, consistent with Huxley-Hill but counter to chemical Fx models, “force maintenance” requires regeneration of an internal mechanical strain, Δµ_ext_ *= F·d*, every time a motor goes around its cycle, and that this internal strain diminishes the chemical energy available for Fx work (left side of Eq. 5). Hill also understood that, consistent with chemical Fx models but counter to Huxley-Hill, the chemistry of force generation is not mechanically isolated from the external force, *F* (i.e., the left side of Eq. 5 is *F*-dependent). Finally, Hill understood that, consistent with Huxley-Hill but counter to chemical Fx models, Fx-work does not influence motor chemistry (it does not appear on the left side of Eq. 5); rather Fx-work is dissipative work that is lost from the motor system to the outside world (it appears on the right side of Eq. 5) through a process made irreversible by the associated heat, *q*, loss.

Because most models of motors and muscle are based on either chemical-Fx or Huxley-Hill chemical-strain formalisms, and a collective force analysis is seldom if ever employed, the conclusions herein suggest that a broad reassessment of the basic mechanisms of muscle and molecular motor function is needed. Our early attempts at simulating collective force generation in muscle yield complex emergent mechanical behaviors that we believe will fundamentally alter our understanding of muscle function in normal and disease states.

## Acknowledgments

This work was supported by a grant 1R01HL090938-01 from the National Institutes of Health.

## References

1. J. Howard, Mechanics of motor proteins and the cytoskeleton (2001; http://www.phy.ohiou.edu/~neiman/biophys-2011/chap8_2.pdf).

2. K. Svoboda, C. F. Schmidt, B. J. Schnapp, S. M. Block, Direct observation of kinesin stepping by optical trapping interferometry. Nature. 365, 721–727 (1993).

3. J. T. Finer, R. M. Simmons, J. A. Spudich, others, Single myosin molecule mechanics: piconewton forces and nanometre steps. Nature. 368, 113–119 (1994).

4. R. W. Lymn, E. W. Taylor, Mechanism of adenosine triphosphate hydrolysis by actomyosin. Biochemistry. 10, 4617–4624 (1971).

5. J. E. Baker, I. Brust-Mascher, S. Ramachandran, L. E. LaConte, D. D. Thomas, A large and distinct rotation of the myosin light chain domain occurs upon muscle contraction. Proc. Natl. Acad. Sci. U. S. A. 95, 2944–9 (1998).

6. A. F. Huxley, Muscle structure and theories of contraction. Prog. Biophys. Biophys. Chem. 7, 255–318 (1957).

7. T. L. Hill, Theoretical formalism for the sliding filament model of contraction of striated muscle. Part I. Prog. Biophys. Mol. Biol. 28, 267–340 (1974).

8. A. F. Huxley, R. M. Simmons, Proposed mechanism of force generation in striated muscle. Nature. 233, 533–538 (1971).

9. M. E. Fisher, A. B. Kolomeisky, The force exerted by a molecular motor. Proc. Natl. Acad. Sci. U. S. A. 96, 6597–602 (1999).

10. H. Qian, A simple theory of motor protein kinetics and energetics. Biophys. Chem. 67, 263–7 (1997).

11. A. B. Kolomeisky, M. E. Fisher, Molecular motors: a theorist’s perspective. Annu. Rev. Phys. Chem. 58, 675–695 (2007).

12. A. D. D. Mehta et al., Myosin-V is a processive actin-based motor. Nature. 400, 590–596 (1999).

13. K. Svoboda, S. M. Block, Force and velocity measured for single kinesin molecules. Cell. 77, 773–84 (1994).

14. J. E. Baker et al., Myosin V processivity: multiple kinetic pathways for head-to-head coordination. Proc. Natl. Acad. Sci. U. S. A. 101, 5542–6 (2004).

15. K. Visscher, M. J. Schnitzer, S. M. Block, Single kinesin molecules studied with a molecular force clamp. Nature. 400, 184–9 (1999).

16. A. Ashkin, Forces of a Single-Beam Gradient Laser Trap on a Dielectric Sphere in the Ray Optics Regime1. Methods Cell Biol. 55, 1–27 (1997).

17. K. Visscher, S. Block, Versatile optical traps with feedback control. Methods Enzymol. (1998) (available at http://www.sciencedirect.com/science/article/pii/S0076687998980405).

18. J. E. Baker, D. D. Thomas, A thermodynamic muscle model and a chemical basis for A. V. Hill’s muscle equation. Mol. Physiol., 335–344 (2000).

19. T. L. Hill, Free Energy Transduction and Biochemical Cycle Kinetics (Springer-Verlag, New York, 1989).

20. T. J. Stewart et al., Actin Sliding Velocities are Influenced by the Driving Forces of Actin-Myosin Binding. Cell. Mol. Bioeng. (2013), doi:10.1007/s12195-013-0274-y.

21. S. Walcott, D. M. Warshaw, E. P. Debold, Mechanical coupling between myosin molecules causes differences between ensemble and single-molecule measurements. Biophys. J. (2012), doi:10.1016/j.bpj.2012.06.031.

22. S. G. Campbell, P. C. Hatfield, K. S. Campbell, A mathematical model of muscle containing heterogeneous half-sarcomeres exhibits residual force enhancement. PLoS Comput. Biol. (2011), doi:10.1371/journal.pcbi.1002156.

23. M. Kaya, Y. Tani, T. Washio, T. Hisada, H. Higuchi, Coordinated force generation of skeletal myosins in myofilaments through motor coupling. Nat. Commun. (2017), doi:10.1038/ncomms16036.

24. J. E. Baker, C. Brosseau, P. B. Joel, D. M. Warshaw, The biochemical kinetics underlying actin movement generated by one and many skeletal muscle myosin molecules. Biophys. J. 82, 2134–47 (2002).

25. L. Gardini, A. Tempestini, F. S. Pavone, M. Capitanio, in Methods in Molecular Biology (2018).

26. R. K. Brizendine et al., Velocities of unloaded muscle filaments are not limited by drag forces imposed by myosin cross-bridges. Proc. Natl. Acad. Sci., 201510241 (2015).

27. A. M. Hooft, E. J. Maki, K. K. Cox, J. E. Baker, An accelerated state of myosin-based actin motility. Biochemistry. 46, 3513–20 (2007).

28. A. V. Hill, The heat of shortening and the dynamic constants of muscle. Proc. R. Soc. London. Ser. B, …. 126, 136–195 (1938).

